# Sibe: a computation tool to apply protein sequence statistics to folding and design

**DOI:** 10.1101/380576

**Authors:** Ngaam J. Cheung, Wookyung Yu

## Abstract

Statistical analysis plays a significant role in both protein sequences and structures, expanding in recent years from the studies of co-evolution guided single-site mutations to protein folding *in silico*. Here we describe a computational tool, termed Sibe, with a particular focus on protein sequence analysis, folding and design. Since Sibe has various easy-interface modules, expressive architecture and extensible codes, it is powerful in statistically analyzing sequence data and building energetic potentials in boosting both protein folding and design. In this study, Sibe is used to capture positionally conserved couplings between pairwise amino acids and help rational protein design, in which the pairwise couplings are filtered according to the relative entropy computed from the positional conservations and grouped into several ‘blocks’. A human *β*_2_-adrenergic receptor (β_2_AR) was used to demonstrated that those ‘blocks’ could contribute rational design at functional residues. In addition, Sibe provides protein folding modules based on both the positionally conserved couplings and well-established statistical potentials. Sibe provides various easy to use command-line interfaces in C++ and/or Python. Sibe was developed for compatibility with the ‘big data’ era, and it primarily focuses on protein sequence analysis, *in silico* folding and design, but it is also applicable to extend for other modeling and predictions of experimental measurements.

---

A protein’s biological function and structure are evolved properties. Co-evolution of amino acids plays a fundamental role in protein evolutionary theory (1). Co-evolution refers to the coordinated changes that occur in pairs of biomolecules or residues to maintain or refine functional interactions between those pairs. To better understand what co-evolutionary information encoded in protein sequences is necessary for protein folding or design, several statistical approaches have been developed in the past decades (2–9). For example, the remarkable results have demonstrated that statistical coupling analysis (SCA) can provide sparsely coupled information for protein design (10), while direct coupling analysis (DCA) is applied to predict residue pair contacts from evolutionary residue-residue couplings (8, 11). Recent studies have shown that existing statistical methods are powerful enough to capture couplings among amino acids for designing foldable proteins (10) and predicting residue contacts (8, 11).

Our ability to reliably detect co-evolutionary information can benefit from the development of additional systematic and well-packed tools that can efficiently and rapidly extract evolutionary information from protein sequences. In computational biology, statistical approaches have been applied to study effects of mutations (variations) on phenotypes, but it is expensive and challenging to design functional assays (12) and create a ‘new’ protein (such as several WW-domain proteins have been created by SCA method and experimentally demonstrated (10)). Protein design has been a long-standing challenge to test the computational approaches in protein sequence analysis, folding and structure prediction, and there have been a number of computational protein design methods (13–19), e.g. creating idealized proteins composed of canonical structural elements (20), the design of coiled coils, repeat proteins, TIM barrels, and Rossman folds (13–18). Moreover, computational results on sequences provided by statistical analysis methods contribute to bridging the gap between protein sequence and design, which may be able to be reduced or filled if the approaches can contribute to protein stability and foldability. Until recently, most of the statistical methods are focused on evolutionary sequence conservation analysis and predictions of residue-residue contacts, which usually are used as constraints for protein structure prediction (11, 21), from the sequences.

Here we investigate whether and how positionally conserved couplings inferred from sufficiently large and diverse multiple sequence alignment (MSA) can be used for specifying a protein’s structure and function, and even to build a ‘better’ version of the protein. We describe the development of a general framework Sibe that meet these challenges. Sibe provides an easy and rapid way for protein design from analytical and computational results on protein sequences. Moreover, we attempt to apply the positionally conserved couplings estimated by co-evolutionary statistics from a protein MSA to define the sequence rules for computationally creating artificial proteins and *in silico* predicting folding pathways & tertiary structures. Two instructive examples are used to show the capabilities of Sibe on protein sequence analysis, design and folding. In the first example, we apply Sibe to statistically analyze an MSA of a eukaryotic signal transduction protein, a G-protein coupled receptor (GPCR) (22), and its design. We use Sibe to build a mutated GPCR protein based on co-evolutionary information inferred from the MSA and compare the functional residues to those in ref. (22). Since the expressive architecture of Sibe, a variety of modules are available to promote its use in both protein engineering and design. In the second example, Sibe is applied to the folding simulation of a group of three proteins based on statistical potentials and the positionally conserved couplings.

## SIBE

Sibe is an analytical and computational framework with functional modules that provides powerful tools for biological science based on modern advances in the field, such as sequence data analysis, *in silico* protein folding and design. All modules in Sibe are implemented in the C++ programming language. It employs an extensible application programming interface (API) in both C++ and Python. Since the introduction of statistical analysis on proteins to the biophysics community, algorithmic improvements for inferring couplings between pairwise residues has been the focus of intense study. Although the practical implementation of these algorithms has presented several historical packages and each of them was strongly tied to the best practice in basic research on protein sequence analysis and folding, software rewrites were common due to the fast-moving research.

Sibe is orientated for extracting meaningful information hidden behind ‘big data’ based on statistical analysis (10, 12) and machine learning (23). With the help of statistical analysis methods, Sibe can infer co-evolutionary information encoded in protein amino acids sequences for folding and design. Through two instructive applications, as illustrated in Fig. 1, we show the capabilities of Sibe. In the first, we apply Sibe to analyze protein sequences for study the relationship between sequence and its thermostability and the analysis could contribute to protein rational design. In the application, Sibe builds a statistical potential of positionally conserved couplings (PCCs) among residues that are accelerated by graphics processing unit (GPU) computing. Due to rapid advances in protein sequence analysis, a variety of APIs is available to researchers. In the second application, we instruct how Sibe implement protein folding using PCCs as drivers and biases that are predicted by machine learning.

**Figure 1:**
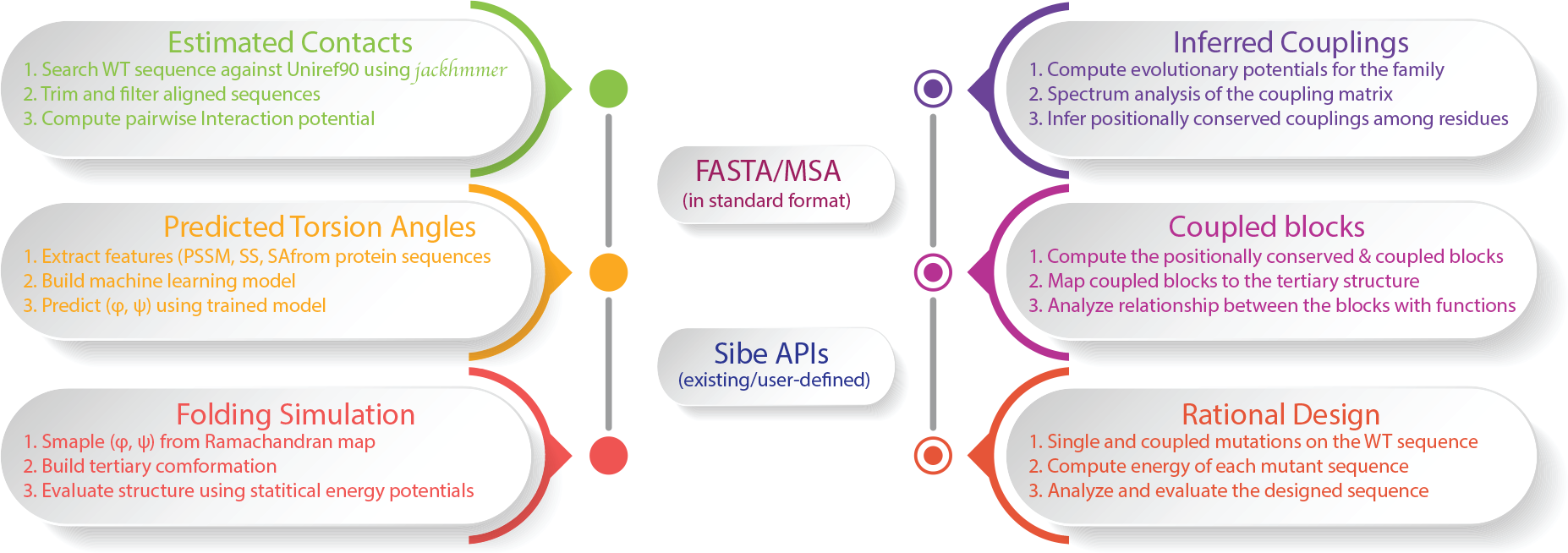
The flowchart of *in silico* protein folding and design in Sibe using evolutionary couplings as drivers.

Sibe incorporates statistical potentials derived from protein sequences (energy-like co-evolution) and structural (energetic potentials (24)) for protein design and folding, respectively. All of the calculations described in this work were carried out within the Sibe software suite and followed the same basic method. In computational protein design, within Sibe large-scale protein sequences were generated by dead-end elimination (DEE) algorithm (25) according to the statistical (energy-like) potential inferred from MSA. Mutants occur in a wild type protein sequence with the guidance of sequence energy-like potential and are accepted by a Metropolis criterion. While the inferences of residue-contacts are also estimated from the MSA as constraint to aid Sibe in protein folding and structure prediction. Combining with predicted constraints of torsion angles (*ϕ* and *ψ*) by a convolution neural network model (26), we performed iterative folding predictions using a Markov Chain Monte Carlo protocol (27) on a set of three representative proteins.

## FROM SEQUENCE TO DESIGN

Written in efficient C++ code, Sibe allows rapid analysis of large protein sequences and captures the evolutionary information for protein folding, design and structure prediction (as illustrated in Fig. 1). In this section, we will describe how statistical analysis in Sibe (as shown in three steps of right Fig. 1) works for a human *β*_2_-adrenergic receptor (*β*_2_AR) protein, which is critical eukaryotic signal transduction protein that communicates across the lipid bilayer. The human *β*_2_AR protein recognizes and binds diffusible ligands (22). Understanding its sequence evolution can provide insight into protein’s functional activities, and help in the design of new drugs and better therapeutics. Here we apply Sibe to study the human *β*_2_AR protein and demonstrate that Sibe can capture significant positionally conserved couplings and important structural features that have been linked to ligand binding activities.

Broadly, the procedure for launching Sibe is to define a set of protein sequences and then align them for estimating frequencies of variations in the sequence alignment. Before starting the statistical analysis, we must obtain the sequences of a given protein we are interested in, and then conduct analysis on the multiple sequence alignment (MSA) for capturing the co-evolution. Generally, the sequences are the output of searching the query against the UniRef90 database (28). In this instructive example, we show in details from sequence alignment to positionally conserved couplings, then to protein design and folding.

### Statistical analysis on sequences

In this study, Sibe focuses on the co-evolution at the residue level including: (1) positionally conserved couplings; and (2) statistical potential derived from site bias of residue and pairwise couplings of residues. As illustrated in Fig. 2, we describe sequence statistics on an MSA. Given an MSA of *N* sequences by *L* positions, denoted as 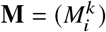, we can obtain an amino acid frequency at an individual position is 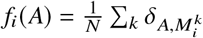, where *δ* = 1 if sequence *k* has amino acid *A* at position *i*, otherwise *δ* = 0. Similarly, a joint frequency of amino acid between a pair of positions is formulated as 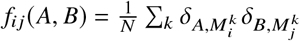.

**Figure 2:**
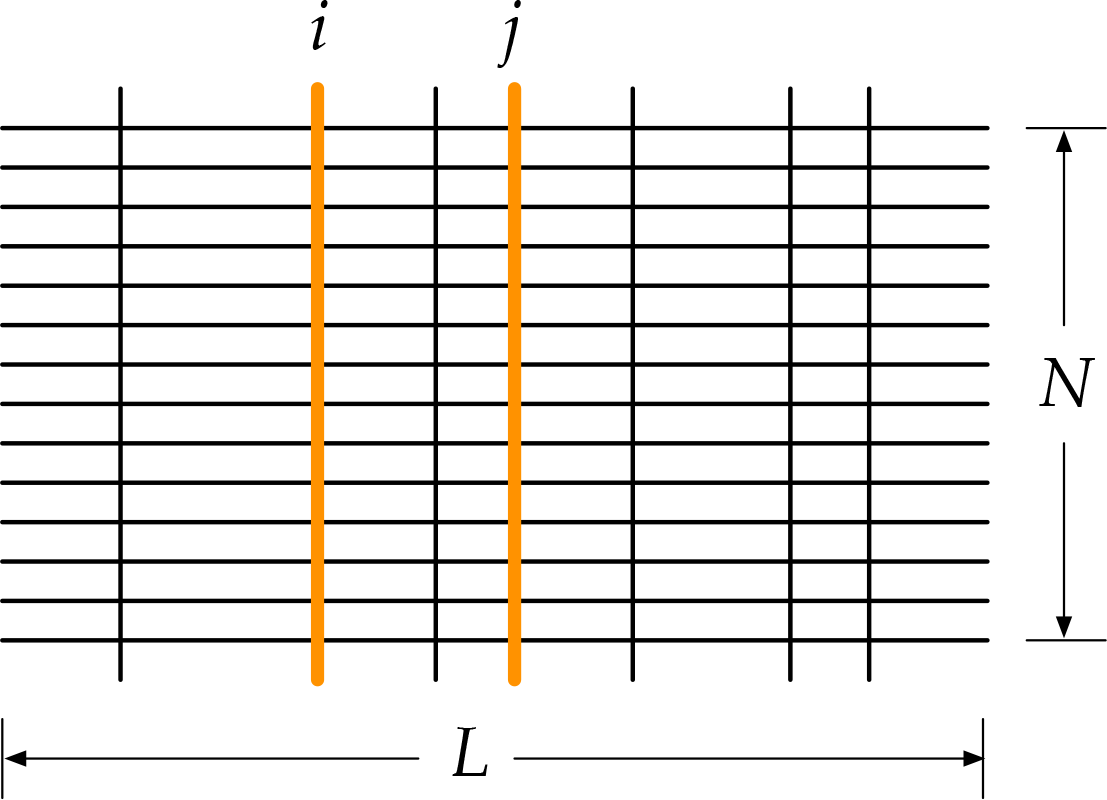
Positionally conserved couplings in a multiple sequence alignment of *N* sequences by *L* positions. The residues at the positions in black lines are conserved, while the residues at positions in bold orange lines are conserved and coupled with each other.

Here, the example of the human *β*_2_AR protein is used to show how Sibe can capture the couplings among residues and an energy-like potential derived from site bias of residue and pairwise couplings of residues. Firstly, we searched the sequence of the *β*_2_AR protein against the UniRef90 database (28) and obtained more than 270,000 sequences. Then we launch Sibe to analyze the MSA of the human *β*_2_AR protein, and we also demonstrate how the statistical energetic potentials derived from the MSA can be used as evolutionary constraints for protein design.

To capture couplings between pairs of residues, we employ the Kullback-Leibler relative entropy (29) to measure how different the observed amino acid *A* at position *i* would be if *A* randomly occurred with an expected probability distribution (4). The definition of the relative entropy is presented as follows,

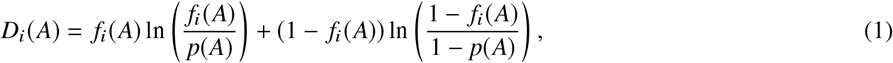

where *p*(⋅) is the background probability.

As shown in ref. (10), the information calculated from the relative entropy of the MSA can remarkably reduce the potential complexity of the protein-design problem. As illustrated in Fig. 3, an example of calculation of the Kullback-Leibler relative entropy from the MSA of the human *β*_2_AR protein. Fig. 3(B) shows a matrix representation of the relative entropy values for twenty different site-specific perturbations calculated from the MSA of the human *β*_2_AR protein.

**Figure 3:**
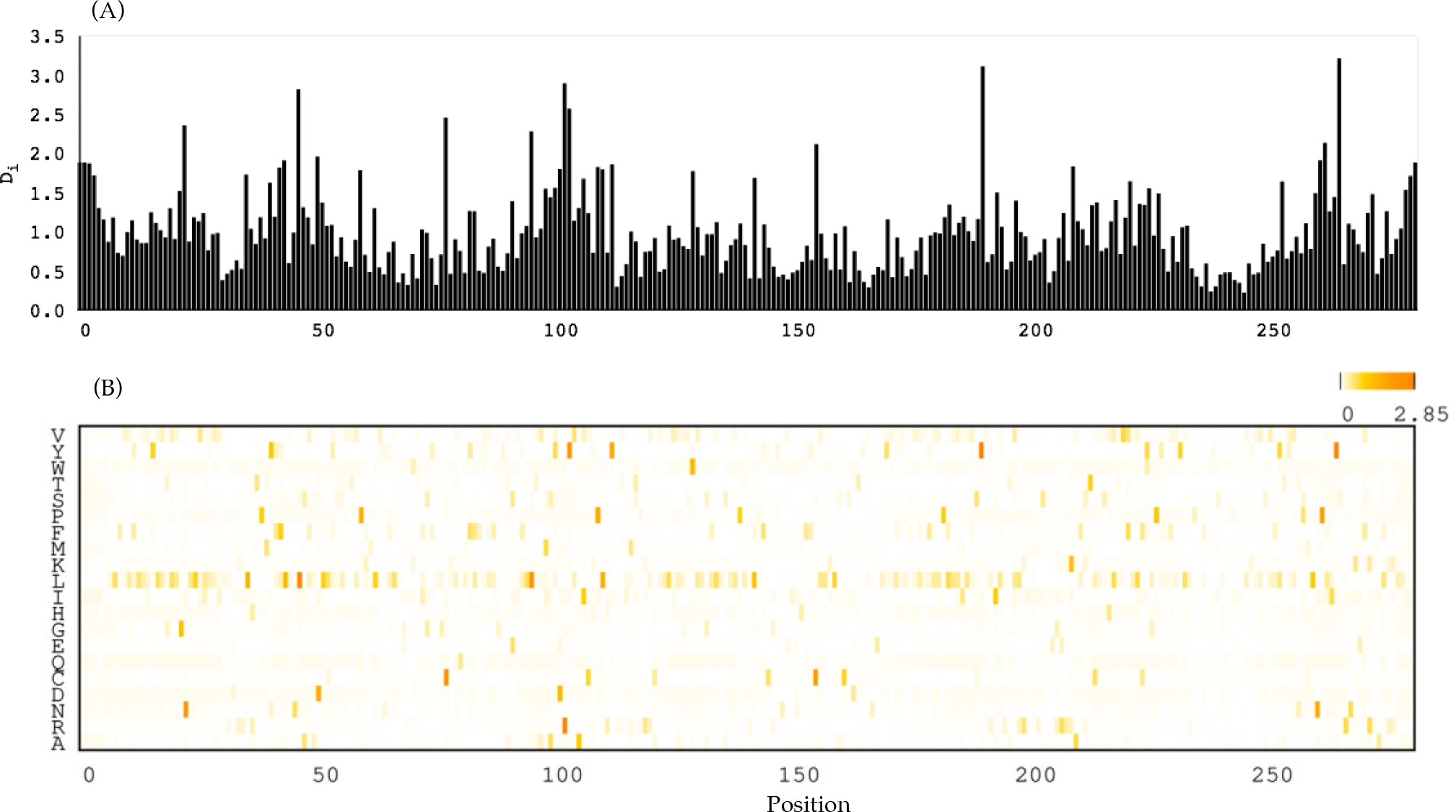
Positional conservation of and Point mutation effect on the human *β*_2_AR protein. (A) Positional conservation in the MSA of the human *β*_2_AR protein. (B) Point mutation effect calculated from the MSA on the human *β*_2_AR protein according to the Kullback-Leibler relative entropy *D*_*i*_. Scale bar shows the relative entropy ranging from 0 (white) to 2.85 (orange).

To capture partial interactions, a global statistical model (DCA-based (5, 6, 8, 11)) is used to infer direct interaction information between pairwise residues. Here, we describe within Sibe how to capture the direct couplings from the given MSA using the model and create an energy-like potential for designing a variant of the human *β*_2_AR protein. Given the MSA, we can easily compute the single site frequency *f*_*i*_(*A*_*i*_) and joint frequency *f*_*ij*_(*A*_*i*_, *A*_*j*_). To maximize the entropy of the observed probabilities, we can calculate the effective pair couplings and single site bias to meet the maximal agreement between the distribution of expected frequencies and the probability model of actually observed frequencies.

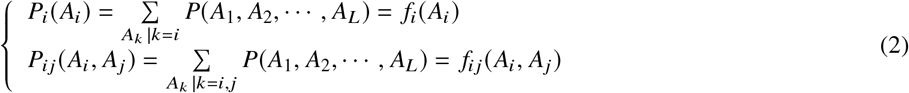

Maximizing the entropy of the probability model, we can get the statistical model as follows,

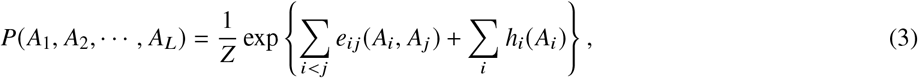

where *Z* is a normalization constant, *e*_*ij*_(·,·), is a pairwise coupling, and *h*_*i*_(·) is a single site bias.

Recently, the DCA-based methods have been reported to be correlated with protein stability and can help design targeted single-point mutation (12, 30, 31). However, since it is difficult to completely disentangle the direct and indirect couplings, they are not always reliable to guide rational protein design. Here, we suggested that positionally conserved couplings between pairwise residues preserve much co-evolutionary information resulting in higher reliability in rational protein design. Rooting in Eqs. (1) and (3), Sibe captures the positionally conserved couplings among residues from the MSA, which contribute to evolutionary constraints for both protein folding and design. In the following paragraphs, we demonstrate that the statistically inferred information of basic evolutionary principles, such as positional conservations and coupled-evolution, can be used to predict protein structures and rationally design a diverse set of ‘better’ proteins.

### Protein sequence design

A great testing ground for the sequence to structure paradigm is protein design (32). However, it is challenging to computationally assay for function in such a large sequence space (12) (e.g. a protein of 25 amino acids has a space of 20^25^). How can we explore the large sequence space to capture key mutations that relate to functional roles of a protein? How can evolutionary information conserved and coupled in the sequences contribute to protein evolution (e.g. kinetic and thermodynamic stability (33)) and lead us to design proteins of novel functional capabilities?

To address the questions, we applied Sibe to facilitate the design of proteins and attempt to uncover the biophysical rules that govern protein folding. In this section, we will use the human *β*_2_AR protein (22) to illustrate how Sibe functions for protein design *in silico* from sequence analysis. In Fig. 4, we provide an overview of the methodology for employing evolutionary couplings as statistical energy-like potential (estimated from a MSA of a protein family by regularized maximum pseudo-likelihood (34)) as constraints for underlying protein sequence design (see also Supplementary Methods).

**Figure 4:**
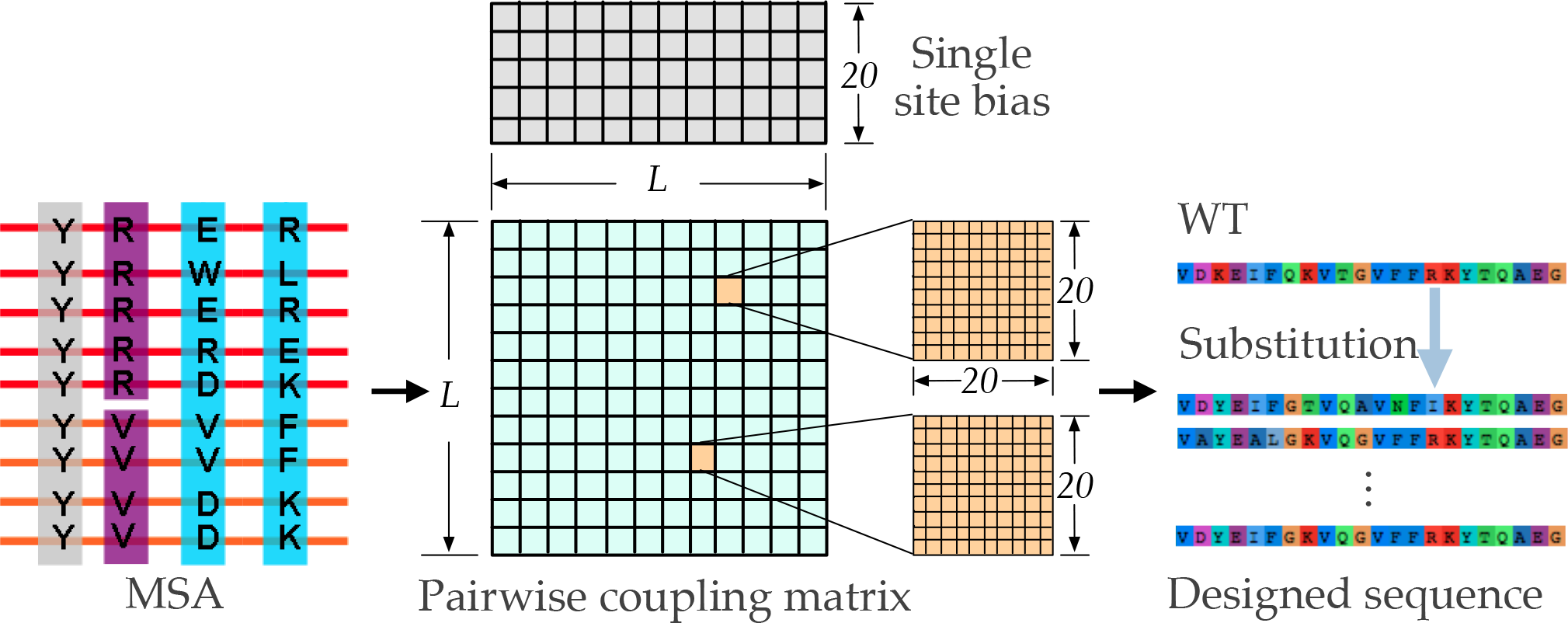
Computational protein design protocol. Energy-like potential is estimated from the MSA, and the DEE is used to *in silico* design the sequences based on the potential.

Firstly, we assessed the energy is found significant correlations with transition temperatures as measured by differential scanning calorimetry experiments for extant and ancestral Trx proteins in study (35), as shown in Fig. 5(A). We searched each given protein sequence as listed in the ref. (35) against the UniRef90 database (28) by HMMER (36) to prepare an MSA for its protein family. The obtained MSA of the given sequence was used as input to create the site bias and coupling matrices. Accordingly, we calculated energies of site bias and couplings for the residues on the sequences. To enhance the ability of the method in distinguishing proteins, we defined an energy equation *E* = *E*_*s*_ + *α* · *E*_*c*_, where *E*_*s*_, *E*_*c*_ and *α* are site bias energy (contribution of a single amino acid to the whole sequence depending on the statistical potential), coupling energy (contribution of pairwise amino acids) and a weight factor, respectively. According to the stability analysis on the thioredoxin family (15 proteins) in the ref. (35), we maximized the correlation between the total energies *E* and the transition temperatures temperatures by optimizing the weight factor (*α*). Based on the calculation, we got a maximum correlation factor about −0.74 when the optimized *α* equals −0.43 as shown in Fig. 5(A).

**Figure 5:**
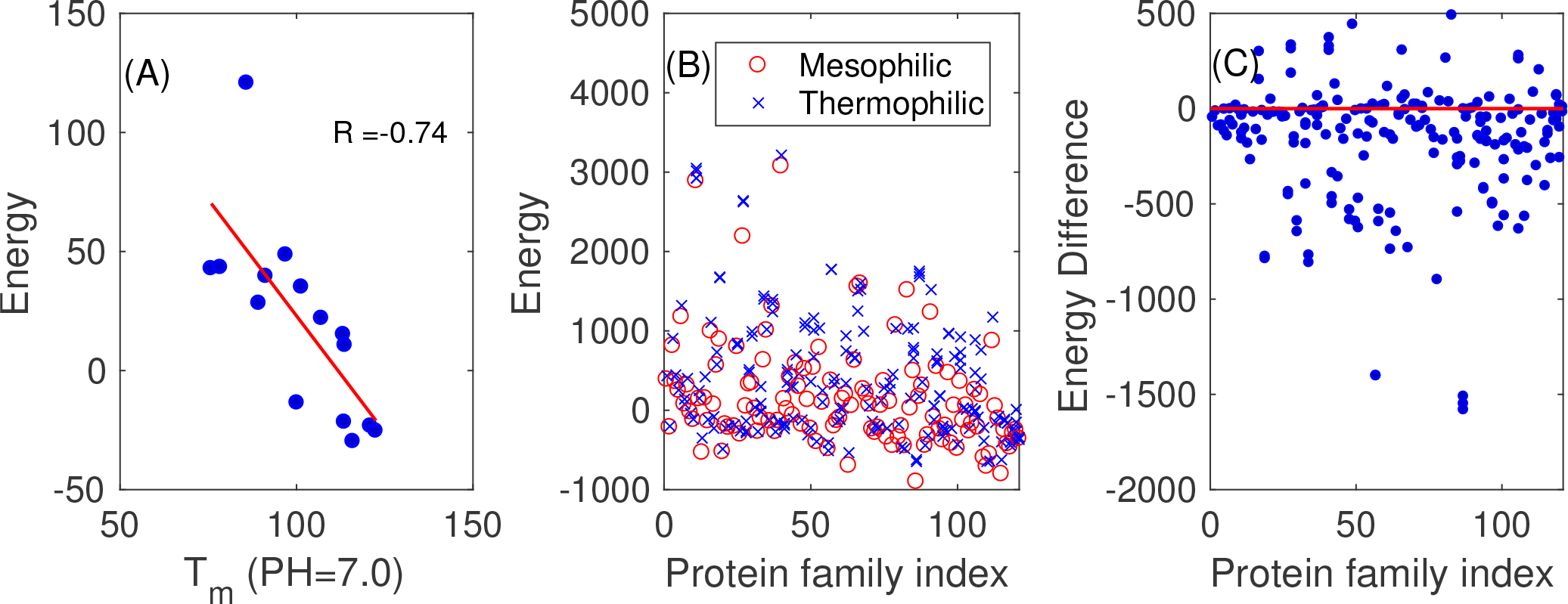
Comparisons between experimental and calculation results. (A) Correlation between the sequence energy and the transition temperatures for extant and ancestral Trx proteins at pH 7.0. (B) The sequence energies of the mesophilic and thermophilic proteins in the different protein families. (C) The energy difference calculated by Sibe between the mesophilic and thermophilic proteins in the same family.

Further, to demonstrate the ability of the derived potential, we applied it to distinguish the mesophilic proteins from thermophilic ones in the same family (see supplementary Table S1). Likewise, we got MSAs of 136 different protein families including mesophilic, thermophilic, moderately thermophilic and extremely thermophilic, and then we launched Sibe to infer the site bias and couplings matrices for the test proteins from the different 136 families. The calculated *E* can distinguish the proteins in the same family and it succeeded on about 83.3% families of all the protein families. As illustrated in Fig. 5(B), red circle and blue cross indicate the energies of thermophilic and mesophilic proteins calculated from the potential in Sibe, respectively. The blue circles in Fig. 5(C) show the differences between the energies *E* of the mesophilic and thermophilic proteins in the same families. As demonstrated on the two experimental data sets, Sibe is able to distinguish mesophilic proteins from thermophilic ones depending on the sequence energy potential.

We next investigated whether Sibe could be used in protein design using the evolutionary information inherent in an MSA. We conducted a computational design study of the human *β*_2_AR protein with the goal of capturing the co-evolutionary information encoding in the natural evolutionary process for designing a new stable variant, which is likely to be functional (to be validated *in vitro*). Accordingly, we calculated the changes in energy for point mutations (Fig. 6), and found that potential mutants are rare (i.e., high energy) in the regions between residues Phe108-Lys149, Phe193-Arg228, His269-Val295, and Tyr313-Leu342 (marked on top of Fig. 6), as well as regions near the carazolol ligand binding sites, residues Trp109, Asp113, Tyr199, Ser203, Trp286, Phe289, Phe290 and Tyr308 (colored in blue at the bottom of Fig. 6) (22).

**Figure 6:**
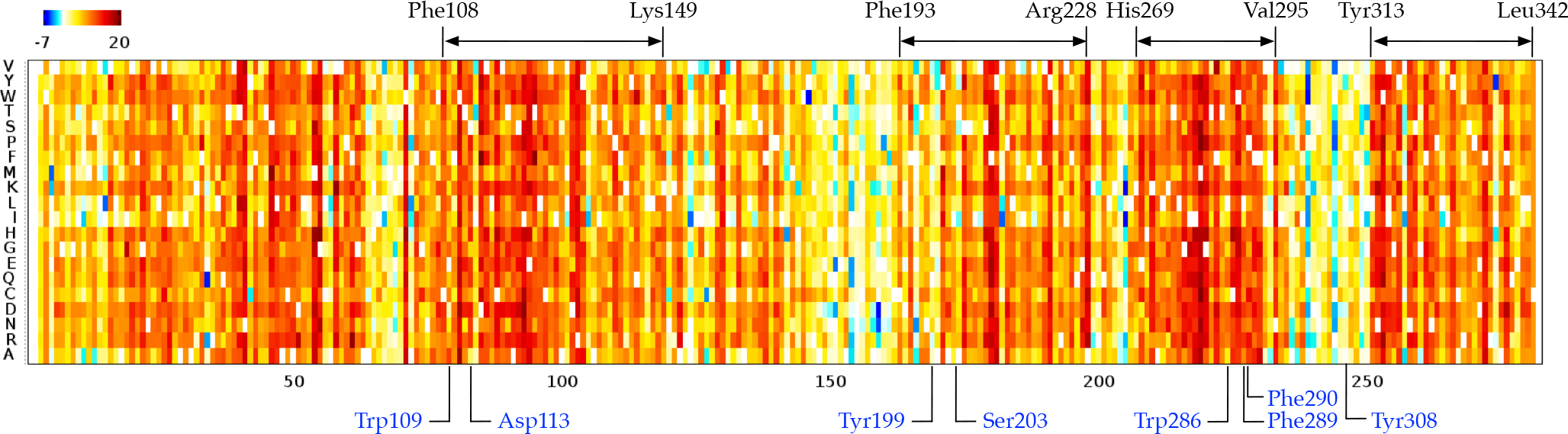
Effects of point mutations on the human *β*_2_AR protein. Substitutions at each position with negative Δ*E* values are predicted to be deleterious; while those that are positive are predicted to be tolerated. Neutral substitutions are marked in 0.

Starting from the estimated potential, we also analyzed the matrix consisting of coupling terms *e*_*ij*_ in Eq. (3) between pair of amino acids by independent component analysis (37). According to the analysis, we found that just the top two eigenvalues are statistically significant. These two top eigenmodes of the coupling matrix (*e*_*ij*_ as illustrated in Fig. 7A) that indicates all the coupled interactions between pairwise residues) are transformed into two independent components for defining independent ‘blocks’ including groups of amino acids. Fig. 7B, C) shows that Sibe successfully obtained two ‘blocks’ for the human *β*_2_AR protein. Each block indicates a group of amino acids that are physically connected (part of the residues in green block are physically connected to each other) in the tertiary structure and may be functionally correlated. It is worth noting that the big red block consists of 39 residues covering functional sites of the protein (22). In principle, the results suggest that Sibe could capture coupled evolution among function-related residues and may contribute to design stabilizing proteins by mutating the wild type residues according to the co-evolutionary information.

**Figure 7:**
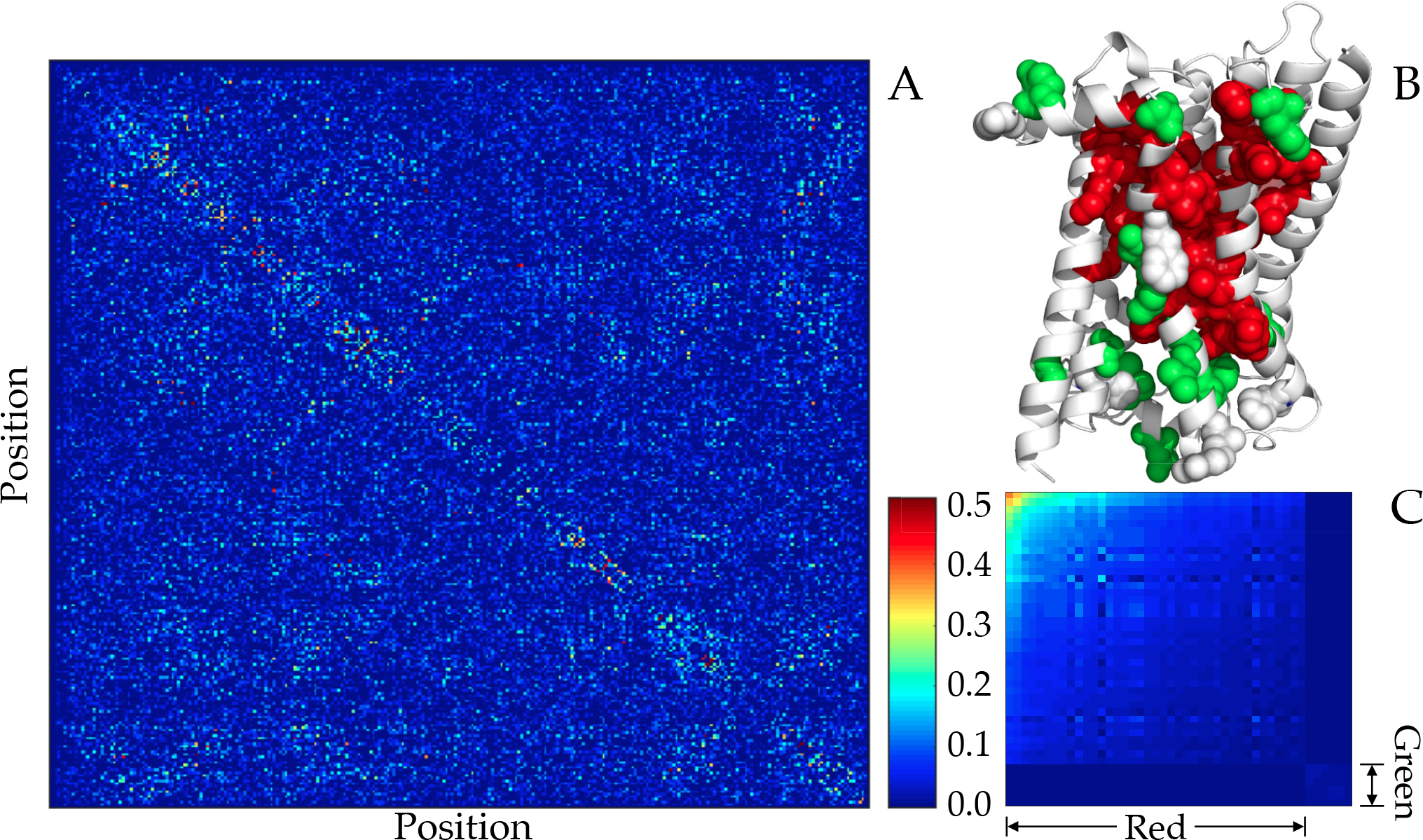
Matrix of pairwise residue-interactions for the human *β*_2_AR protein. (A) Evolutionary interactions inferred by PCC between pairwise residues. (B) Two interaction ‘blocks’ mapped to the tertiary structure. (C) Coupling matrix reordered to highlight two interaction blocks. Those blocks show that the residues in the same block may contribute similarly functions to the protein.

The critical feature of the design protocol is described in supplementary materials. Sibe can be used to identify of the designed sequence with the lowest energy. To assess whether the approach can produce a ‘better’ protein from coupling constraints encoded in the MSA, according to the co-evolution derived energies, we performed the DEE minimization protocol (25) on sequence design. For each starting sequence, five thousand independent simulations, each trial of sequence design with maximum iterations of 100,000, were performed to obtain a sequence of the lowest energy. As illustrated in Fig. 8, we show the ligand-binding characterization in the human *β*_2_AR protein. As shown, the residues colored in green in Fig. 8 are the ligand-binding sites, where extensive interactions occur between the human *β*_2_AR protein and carazolol at positions Phe289, Phe290 and Trp286 (22). In the designed human *β*_2_AR protein, we got three mutants in the ligand-binding sites, which may alter the function of the designed protein.

**Figure 8:**
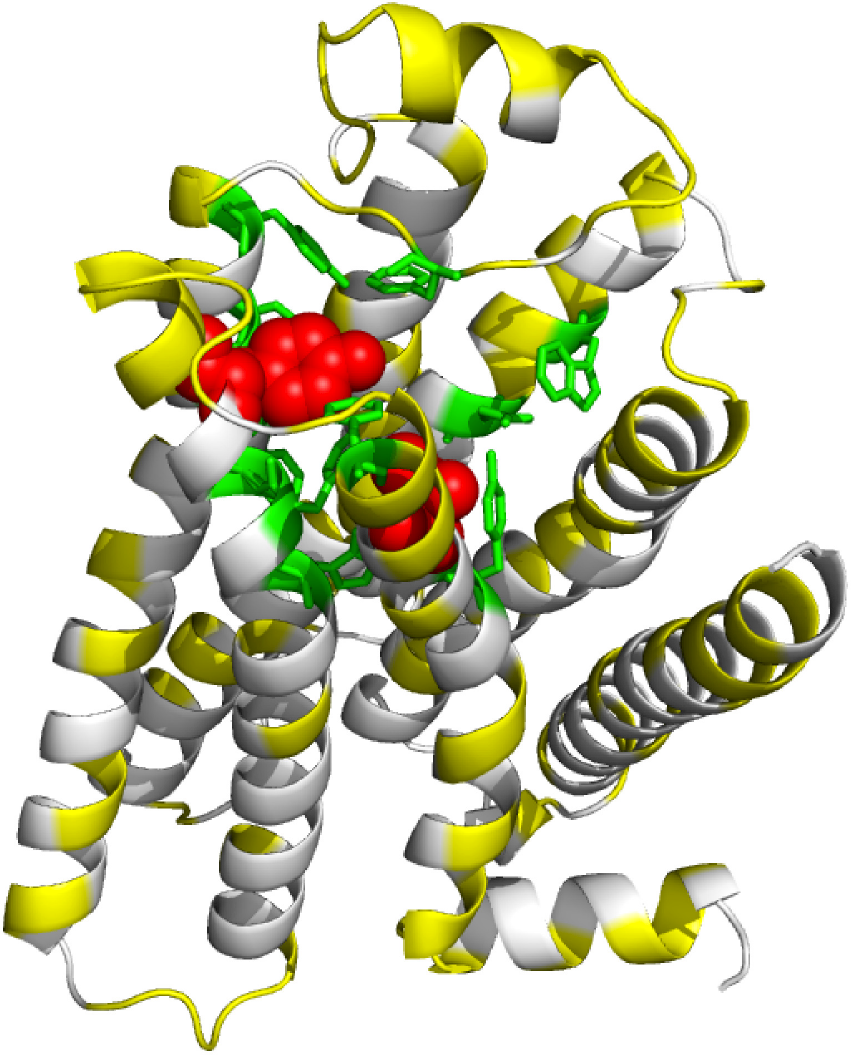
Sibe computationally designed a mutant sequence of the human *β*_2_AR protein (PDB ID: 2RH1) (22) that is mapped to it tertiary structure. Yellow, green and red indicate mutations (made by Sibe) occurred at common sites, ligand-binding sites, and experimentally mutated functional sites, respectively.

## PROTEIN FOLDING AND STRUCTURE PREDICTION

Prediction of protein structure has been a long-standing challenge, and numerous advances have been made in determination of the three-dimensional structure of a protein from its amino acid sequence or primary structure (24, 38–40). Recently, due to efforts in metagenome sequence projects, the number of protein sequences is growing considerably faster than ever before (21, 41). However, there are remaining challenges regarding efficient computational methods for interpretation of large sequence data in protein families and rapid structure modeling approaches bridging the gap between sequences and corresponding structures. The gap between a protein sequence and its unknown structure can be also largely reduced by taking advantage of progress in statistical analysis in both protein sequences and known structures. Moreover, co-evolution among amino acids enhances capacities of existing computational approaches in predictions of contacts between different protein residues (5, 6, 8), and the predicted constraints can provide an accurate way in modeling a protein of unknown structure (21).

To assess structure prediction, we carried out calculations on an instructive example consisting of three representative proteins according to the three steps as shown in the left of Fig. 1, the low-molecular-weight protein tyrosine phosphatases YwlE (PDB ID: 1ZGG), the flagellar capping protein (PDB ID: 5FHY), and the E. coli MCE protein MlaD (PDB ID: 5UW2), using positionally conserved couplings and predicted protein (*ϕ, ψ*) torsion angle constraints (23). We present an iterative framework (Fig. 9) to fold a protein using statistical potentials (24) and co-evolution constraints derived from its sequence alignment (as described above). The iterative prediction using a Markov Chain Monte Carlo protocol (Supplementary Method) includes multiple rounds, in which the predicted constraints (e.g. torsion angles, residue-contacts) can be collected from the previous round to guide and bias the prediction.

**Figure 9:**
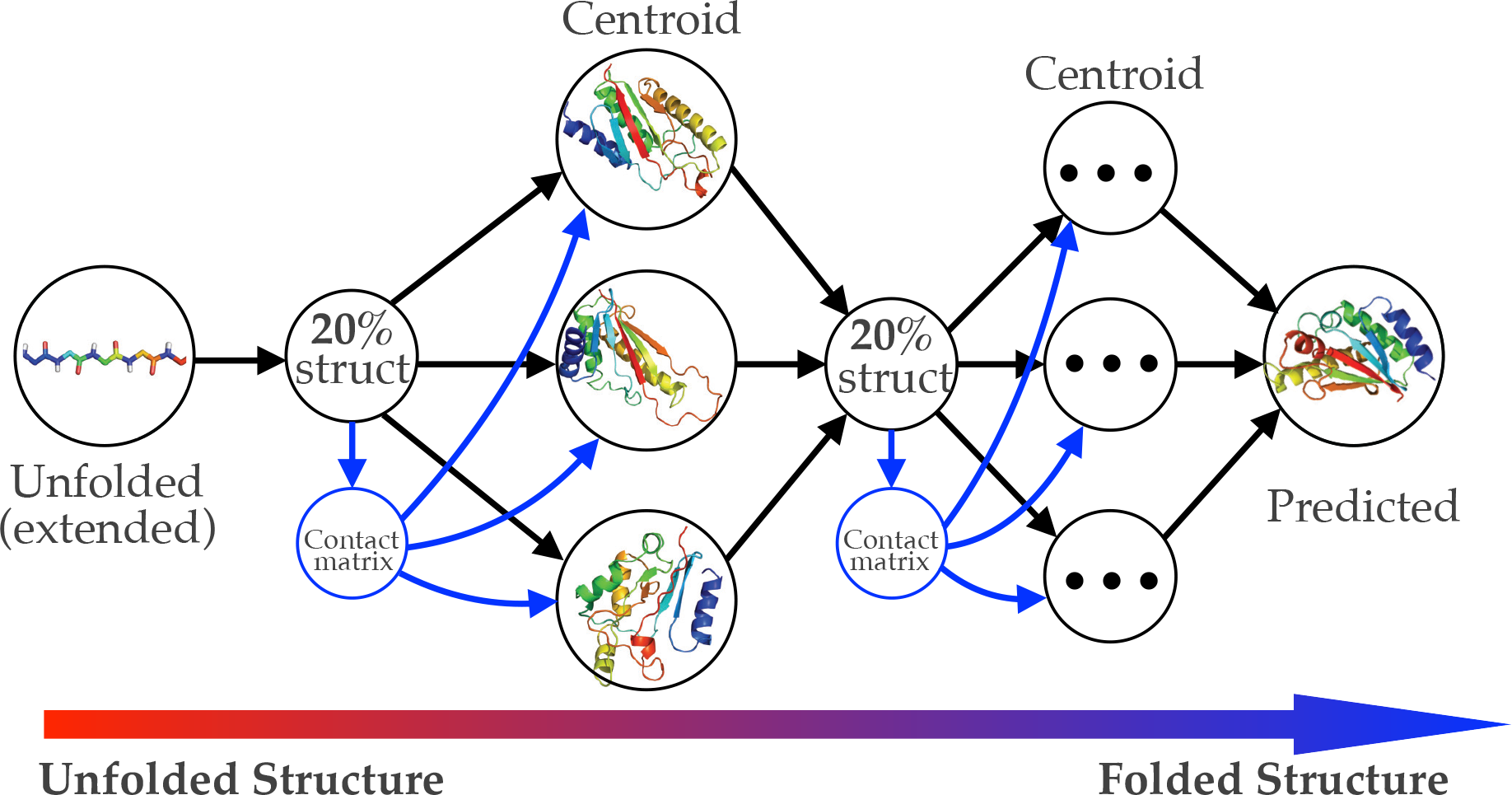
Iterative structure prediction guided by co-evolution. Starting from unfolded (extended) structure, Sibe incorporates residue-contacts derived from coupling analysis on the MSA and averaged residue-contacts from predicted structures (previous round) with lower energy (best 20%) as constraints to iteratively predict the tertiary structures of targets.

Since each potential confirmation of a protein is drawn from the same Ramachandran map distribution according to the given amino acid sequence, the conformations generated by the Markov Chain Monte Carlo method are partially correlated with each other. In the folding simulation, starting from a query sequence, we generated five hundred initial conformations to trigger its structure prediction by iteratively biasing the prediction from constraints. For example, Sibe was used to iteratively predict the tertiary structure of the YwlE protein using the constraints of residue-contacts inferred from its MSA and predicted torsion angles (which are used to increase probabilities of *ϕ* and *ψ* located in 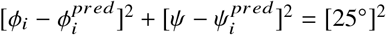 on the Ramachandran map distribution) predicted by *Phsior* (23). After 500 single simulations were converged, we chose a hundred of the predicted structures with the lowest energy (20% of all structures) and calculated the averaged residue-contacts (for spatial interactions among residues) and torsion angles (define the square ranges for each pair of *ϕ* and *ψ*) (see also supplementary methods).

As illustrated in Fig. 10, we present the predicted models for the target proteins by comparison to the crystal structures. The predicted results show the capability of the module of protein structure prediction in Sibe is demonstrated by the blind predictions of protein tertiary structures close to the crystal structures.

**Figure 10:**
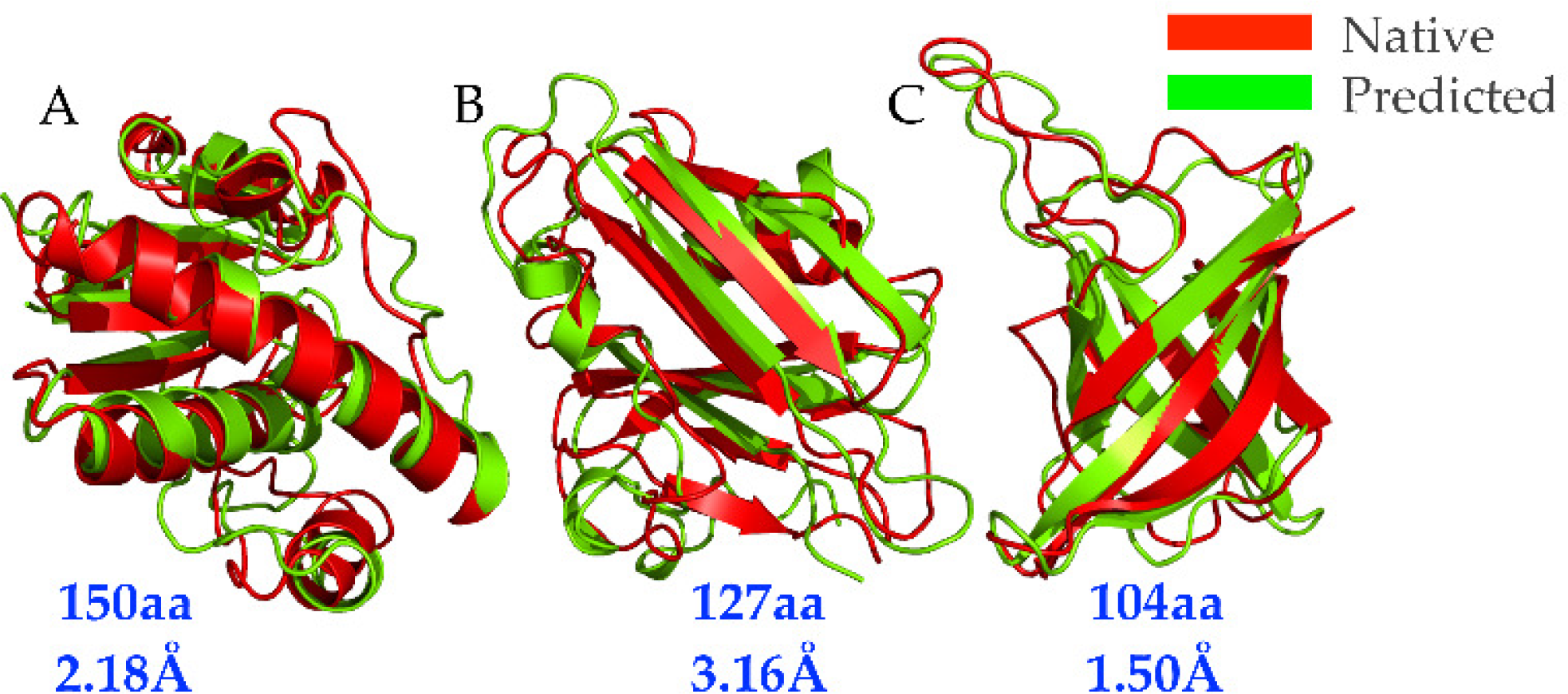
Comparison of models predicted by Sibe (green) to the crystal structures (red). The models accurately recapitulate the structural details of the named proteins. The RMSD of each protein was computed by PyMOL software (42) as shown, and the TM-scores are as follows: (A) YwlE (PDB ID: 1ZGG, TM-score 0.76), (B) the flagellar capping protein (PDB ID: 5FHY, TM-score 0.64), (C) the E. coli MCE protein MlaD (PDB ID: 5UW2, TM-score 0.80).

## CONCLUSION

We report here that a software suite (Sibe) provides an analytical and computational tool for protein folding & design studies. Sibe allows rapid analysis of large protein sequences data for boosting protein design and folding.

The success of the software depends in part on the positionally conserved couplings derived method for detecting amino acid variations, and the easy-interface modules presented in Sibe lay the ground-work for drawing interpretable conclusions from protein sequence data to its folding and design studies *in silico*. For example, due to rapid advances in the software suite Sibe, a variety of functional modules are available to researchers for analyzing protein sequences, protein folding and design *in silico*. In the second example, we demonstrate how Sibes implementation of iteratively biasing conformation search can be used to predict the tertiary structure from its amino acid sequence based on statistical potentials of protein sequences and structures. Due to the limited diversity in the MSA of a given protein family, Sibe is imperfect in capturing significant co-variants as co-evolutionary constraints for protein design and structure prediction. Accordingly, the remaining challenges are how to enrich the diversity information in the MSA and how to efficiently detect important co-evolutionary couplings between pairwise amino acids and distinguish the couplings from biases that arise from a protein family of less diversity. Future work may focus on addressing those issues by extension and improvement of Sibe.

Generally, Sibe’s power and simple architecture are dependent on expressive and functional modules, which focus on extending methods specifically designed for scientific applications. Understanding the co-evolutionary process from metagenome sequence data provides thermodynamic insights into the protein’s evolution, which can aid in the design of better proteins. Hopefully, the methodology of protein design could have future applications in chemistry, bioremediation, and drug design & discovery.

## AUTHOR CONTRIBUTIONS

NJC and WY designed the research. NJC carried out all simulations, together with WY analyzed the data. NJC and WY wrote the article.

## AVAILABILITY

The Sibe software suite can be obtained at request for non-commercial use and its installation documentation is available on its web-server at: http://wyu.dgist.ac.kr/sibe.

## ACKNOWLEDGMENTS

We thank the Freed/Sosnick group members, Drs. S. Wang, and J. Jumper for helpful discussions. This work was supported by the grant (NRF-2017R1D1A1B03031845).

