## Supplemental materials for "Sibe: a computation tool to apply protein sequence statistics to folding and design"

N. J. Cheung, Wookyung Yu

### Methods and materials

Sibe is a software suite rooting in statistical models, convolutional neural network, optimization solvers and energy functions for both protein studies and data mining. In this study, within Sibe statistical information inferred from multiple sequence alignment is generated and applied to protein folding, structure prediction and design. Two main tasks were presented by two instructive examples, in which we are to show the ability of Sibe in folding and design a protein.

#### Analysis on protein sequences

First of all, we search a query sequence against the UniRef90 database [1] by HMMER [2] and then prepare its sequence alignment. The following command lines are used to collect the aligned sequences,

```
jackhmmer \
 --notextw \
  -A out.sto \
 --tblout out_tbl.out \
 --domtblout out_domtbl.out \
  -E 0.01 \
 --popen 0.25 \
 --pextend 0.4 \
   query_fasta \
   uniref90.fasta
```

We prepared a wild type sequence of the human $\beta_2$AR protein as the example for protein sequence analysis. By running `jackhmmer`, we got a raw sequence alignment and then trimmed them by a command line as follows,

```
sibe sequence_trim \
 -msa=raw_sequence_alignment.msa
```

The command line is to trim the aligned sequences according to the first sequence (query sequence) in the input sequence alignment.

At the begin of sequence analysis, we use command `sequence_stats` in Sibe to conduct a naive examination of the sequence space described by the alignment. Sibe will also present a Shell script for plotting figures of the output. By running the script, the computed a matrix of similarity

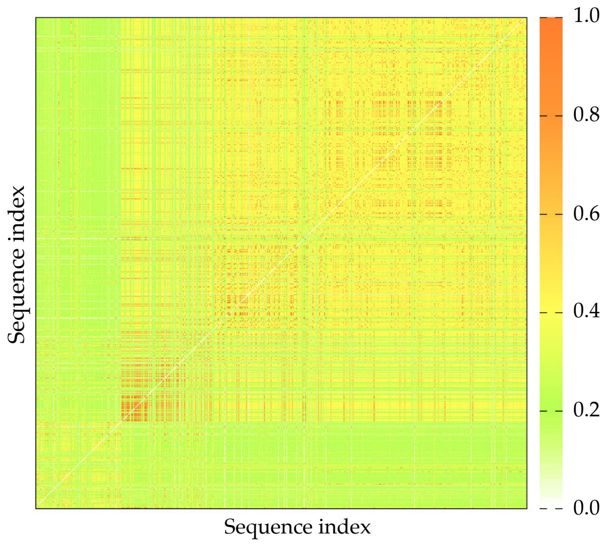


Figure S1: Similarity between pairs of sequences.

between pairs of sequences can be plotted as illustrated in Fig. S1. The matrix representation gives the fraction of amino acids that are common between the pairwise sequences. As another computational results, the bar representation of positional conservation is shown in Fig. 3.

```
sibe sequence_stats \
-msa=raw_sequence_alignment.msa
```

### Computational protein design

#### Validations of the calculations

In computational protein design, we first demonstrate that the estimated sequence potential from MSA can distinguish the mesophilic and thermophilic proteins. Likewise, we prepared a MSA for each family and then inferred the energy-like potential from the MSA by running command line as follows,

```
sibe sequence_coupling \
 -msa=sequence_alignment_trimed.aln
```

Based the obtained potential, we calculated the energies of the sequences using the same estimated potential if the sequences are in the same family. The command line is to compute the sequence energy as follows,

```
sibe sequence_energy \
 -msa=the_sequences_require_energy_calculation.aln \
 -mat=sequence_potential.mat
```

The calculations are shown in Fig. 6. We demonstrated that the computational results have high correlation (Pearson correlation coefficient $\sim -0.74$) with the experimental data [3], and the estimated potential and the model were validated to be efficient in distinguishing the mesophilic from thermophilic proteins, 24 of all protein families as listed in Table S1, others can be find in ref. [4].

Table S1: Meso- & thermo-philic proteins used in this study

| No. | Family name | PDB ID | Source organism (meso-/thermo-philic) |
|---|---|---|---|
| 1 | Transcription initation | 1volA | Human (meso) |
| | factor IIb | 1aisB | Pyrococcus woesei (thermo/100°C) |
| 2 | Superoxide dismutase | 1ar4A | Propionibacterium freudenreichii (meso) |
| | (Mn- or Fe-dependent) | 3mdsA | Thermus thermophilus (thermo/75°C) |
| 3 | Glutamate dehydrogenase | 1hrdA | Clostridium symbiosum (meso) |
| | | 1gtmA | Pyrococcus furiosus (thermo/100°C) |
| 4 | Malate dehydrogenase | 4mdhA | Pig heart (meso) |
| | | 1bmdA | Thermus flavus (thermo/72.5°C) |
| 5 | Phycocyanin alpha chain | 1cpcA | Fremyella diplosiphon (meso) |
| | | 1liaA | Polysiphonia urceolata (meso) |
| | | 1phnA | Cyanidium caldarium (thermo/45°C) |
| 6 | Signal recognition particle | 1fts | Escherichia coli (meso) |
| | (receptor) | 1ffh | Thermus aquaticus (thermo/72.5°C) |
| 7 | Ferredoxin | 1fxd | Desulfovibrio gigas (meso) |
| | | 1fxrA | Desulfovibrio africanus (meso) |
| | | 1vjw | Thermotoga maritima (thermo/80°C) |
| 8 | Subtilisin | 2pkc | Tritirachium album limber (meso) |
| | | 1thm | Thermoactinomyces vulgaris, 60°C) |
| 9 | Neutral protease | 1npc | Bacillus cereus (meso) |
| | (thermolysin) | 1lnfE | Bacillus thermoproteolyticus (thermo/52.5°C) |
| 10 | Rubredoxin | 6rxn | Desulfovibrio desulfuricans (meso) |
| | | 1caa | Pyrococcus furiosus (thermo/100°C) |
| 11 | Cyclodextrin | 1cdg | Bacillus circulans strain 251 (meso) |
| | glycosyltransferase | 1cgt | Bacillus circulans strain 8 (meso) |
| | | 1pamA | Bacillus sp. 1011 (meso) |
| | | 1ciu | Thermoanaerobacterium thermosulfurigenes (thermo/60°C) |
| | | 1cyg | Bacillus stearothermophilus (thermo/52.5°C) |
| 12 | Phycocyanin beta chain | 1cpcB | Fremyella diplosiphon (meso) |
| | | 1liaB | Polysiphonia urceolata (meso) |
| | | 1phnB | Cyanidium caldarium (thermo/45°C) |
| 13 | 3-Phosphoglycerate kinase | 1qpg | Yeast (meso) |
| | | 1vpe | Thermotoga maritima (thermo/80°C) |
| 14 | Glyceraldehyde-3-phosphate | 1a7kA | Leishmania mexicana (meso) |
| | dehydrogenase | 1hdgO | Thermotoga maritima (thermo/80°C) |
| 15 | Xylanase (I) | 1ukrA | Aspergillus niger (meso) |
| | | 1yna | Thermomyces lanuginosus (thermo/45°C) |
| 16 | Xylanase (II) | 1clxA | Pseudomonas fluorescens (meso) |
| | | 1xyzA | Clostridium thermocellum (thermo/60°C) |
| 17 | TATA box binding protein | 1cdwA | Human (meso) |
| | | 1vokA | Arabidopsis thaliana (meso) |
| | | 1pczA | Pyrococcus woesei (thermo/100°C) |
| 18 | Adenylate kinase | 2ak3A | Bovine (meso) |
| | | 1ukz | Yeast (meso) |
| | | 1zip | Bacillus stearothermophilus (thermo/52.5°C) |

| | | | |
|---|---|---|---|
| 19 | Carboxypeptidase | 1nsa | Pig (meso) |
| | | 1pca | Pig (meso) |
| | | 1obr | Thermoactinomyces vulgaris (thermo/55°C) |
| 20 | Ornithine carbamoyltransferase | 2otcA | Escherichia coli (meso) |
| | | 1a1s | Pyrococcus furiosus (thermo/100°C) |
| 21 | Pyrophosphatase | 1obwA | Escherichia coli (meso) |
| | | 2prd | Thermus thermophilus (thermo/72.5°C) |
| 22 | CheY protein | 3chy | Escherichia coli (meso) |
| | | 2chf | Salmonella typhimurium (meso) |
| | | 1tmy | Thermotoga maritima (thermo/80°C) |
| 23 | Phosphofructokinase | 1pfkA | Escherichia coli (meso) |
| | | 4pfk | Bacillus stearothermophilus (thermo/52.5°C) |
| 24 | Triacylglycerol acylhydrolase | 1lgyA | Rhizopus niveus (meso) |
| | | 1tib | Humicola lanuginosa (thermo/50°C) |
| | | 3tgl | Rhizomucor miehei (thermo/45°C |

The effects of point mutations were computed from the potential by running following command line. The matrix representation of the computational results is shown in Fig. 7.

```
sibe point_mutation \
 -fastx=query_fasta_WT.fasta \
 -mat=input_sequence_potential.mat
```

#### Protein design protocol

Using the algorithm described by Desmet et al. [5], the DEE procedure for each trajectory starts with a given wild type sequence. The sequence converges to a local energy minimum by gradually decreasing the temperature. As illustrated in Fig. S2, starting from a given wild type sequence (`WT_sequence`), we launched a DEE algorithm to optimize the mutant sequence base on the energy-like potential (`sequence _potential`) inferred from the MSA. According to the Metropolis criterion, the design protocol will accept or reject a new mutant that may occur in the wild type sequence, and the change is accepted with probability,

$$P = \min\{r_p, e^{-\Delta E/t}\} \tag{1}$$

where $r_p = 1$, and $\Delta E$ is the energy difference between the new and old designed sequences.

By running the following command line, we can get a trajectory (`the_designed_sequence .trajct`) of mutant sequences starting from the wild type sequence.

```
sibe sequence_design \
 -fastx=WT_sequence \
 -mat=sequence_potential.mat \
 -dseq=the_designed_sequence.trajct \
 -iterations=100000
```

From the design protocol, we got five hundreds trajectories of the designed sequences, and the sequence with lowest energy in each trajectory was collected for visualization as shown in Fig. S4.

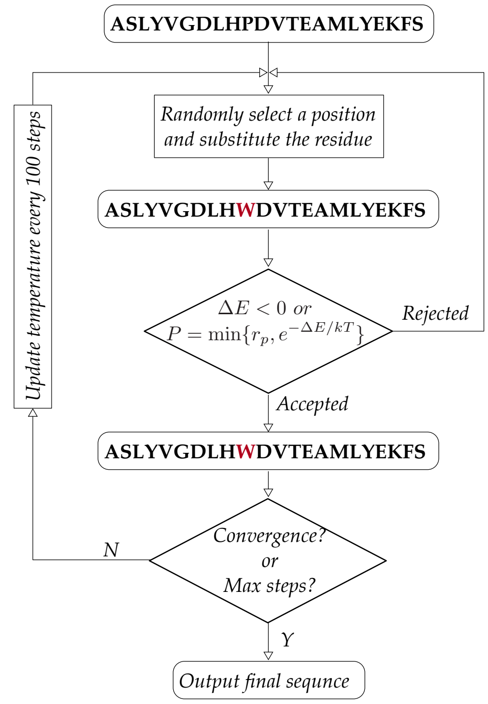


Figure S2: Flowchart of the computational protein design.

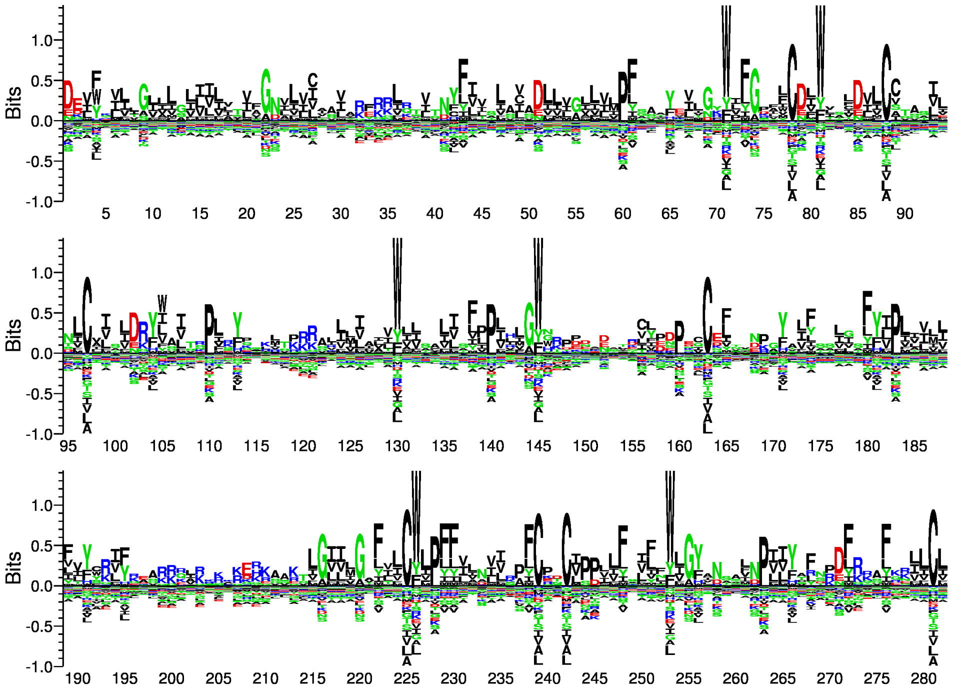


Figure S3: Visualization of the computationally designed sequences with the lowest energy in each trajectory.

### Protein folding and structure prediction

In Sibe, MCMC algorithm is used for protein conformation sampling from individual ($\phi$, $\psi$) distributions (Ramachandran maps), while all other angles and bond lengths are fixed at their ideal values. A single round involves 500 individual MCMC folding simulations that are run using specialized ($\phi$, $\psi$) backbone sampling procedures and the energy functions (as described in ref. [6]) in a protein representation containing the backbone and $C_\beta$ atoms. Within the simulation, MCMC provides a general solution to protein folding and structure prediction prevalent in scientific research. As described in ref. [6], Sibe utilizes the same moving sets and energy functions. Additionally, Sibe employed predicted angles ($\phi$, $\psi$) to increase the sampling probabilities in the Ramachandran map distribution, which also efficiently enhance the ability of the MCMC method during the simulation. The passing of constraints of torsion angles ($\phi$, $\psi$) and residue-contacts from one round to another is repeated until convergence as illustrated in Fig. S4.

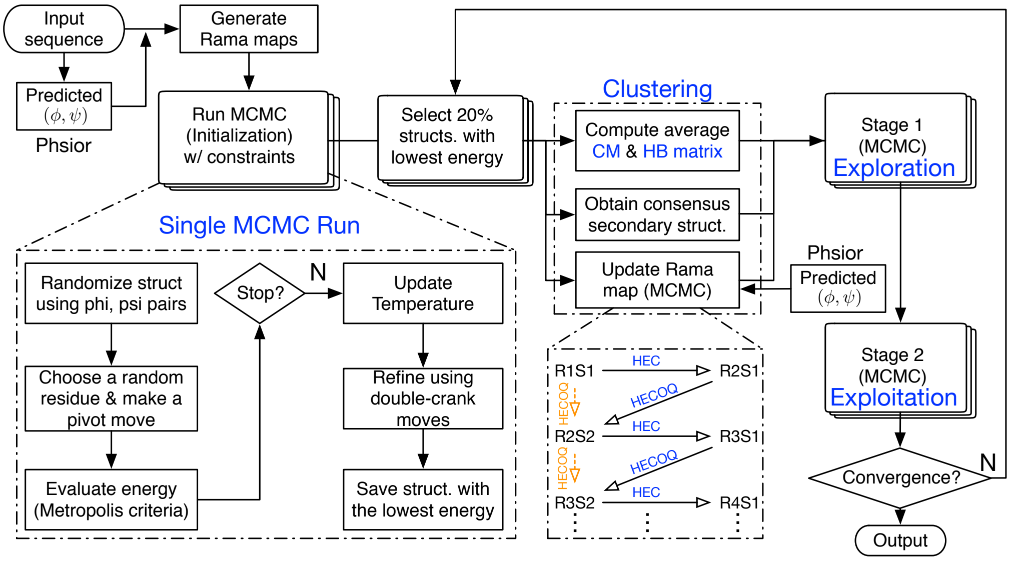


Figure S4: Flowchart of the prediction system.

### Constraints-guided sampling

In Sibe, we used the *Phsior* [7] to predict torsion angles ($\phi$, $\psi$), which were applied to shift the Ramachandran map distribution and increase the probabilities $p$ of ($\phi$, $\psi$) of the $i$th residue located in circle $[\phi_i - \phi_i^{pred}]^2 + [\psi - \psi_i^{pred}]^2 = [25°]^2$ by increment of $0.75 * p$. In each round, a consensus secondary structure is obtained at every amino acid position, which is used to alter the Ramachandran map distribution. Combining with the predicted torsion angles ($\phi$, $\psi$), we can make changes in the probabilities in each map of amino acids. This strategy leads to efficient samplings in the MCMC algorithm and accelerate the folding of a protein structure.

The residue-contacts (estimated by DCA [8, 9]) are used as constraints in iterative simulations as illustrated in Fig. 10. The top L/3, L/2, L, 3L/2 (L is the length of a protein sequence) predicted contacts are used for 500 simulations in initial stage. Every 125 simulations utilize the same initial constraints of residue-contacts, e.g. top L/3 residue-contacts are used in the first 125 simulations while top L/2 residue-contacts are employed by another 125 simulations. This hierarchical application of constraints from residue-contacts is to help the Markov chain sample efficiently in conformation space.

### Metropolis-Hastings sampling

In Sibe, we use a Markov chain to sample from Ramachandran maps distribution ($\pi$) (derived from high resolution PDB structures [6]). Accordingly, it is necessary to develop a transition for the Markov chain, aiming to match the chains stationary distribution with the individual ($\phi$, $\psi$) distributions. As an effective strategy to sample angular space ($\phi$, $\psi$), a random-walk Markov chain uses the current state of a chain of conformations to propose a new state. Within Sibe, a Gaussian function centered on the current state $g(s|s^{(t-1)}) \sim \mathcal{N}(s^{(t-1)}, 1)$ is defined as the proposal function. This allows the algorithms to exploit the conformation space of the posterior–if a new conformation is similar to the last draw, then it is likely to be accepted. The proposal distribution $g(s|s^{(t-1)})$ is dependent only on the previous state in the Markov chain.

Starting from some random initial state $s^{(0)} \sim \pi^{(0)}$, the protocol first draws a potential state $s$ from a proposal distribution $g(s|s^{(t-1)})$. Accordingly, in Sibe the Metropolis-Hastings algorithm is launched as the simplest form of random-walk MCMC protocol, which accepts or rejects a transition from protein conformation $s^{(t-1)}$ to $s$ with probability $\alpha$. The detailed criteria for accepting or rejecting a proposed state are defined as follows:

1. If $R(s) \geq R(s^{(t-1)})$, the proposed state $s$ will be set as the next state in the Markov chain.

2. If $R(s) < R(s^{(t-1)})$, then the proposed state may still be accepted, but only randomly, and with a probability $\frac{R(s)}{R(s^{(t-1)})}$. The acceptance probability for the proposed state is calculated as follows,

$$P(s^{(t-1)} \to s) = \min\left(1, \frac{R(s)g(s \to s^{(t-1)})}{R(s)g(s^{(t-1)} \to s)}\right), \tag{2}$$

   where $R$ is the density of the individual $\phi$, $\psi$ distribution of an amino acid, and $g$ is the density of the proposal function for a transition from $s^{(t-1)}$ to $s$.

   The proposed state $s$ is accepted if a random uniform number $u$ is less than or equal to $\alpha$. Otherwise, it will be rejected, and the current state will be set as the next state in teh Markov chain.
